# YAP activity sustains mitochondrial adaptation to NAMPT inhibition in triple-negative breast cancer

**DOI:** 10.64898/2026.11.18.766653

**Authors:** Arianna Buongiorno, Elia Franceschi, Elena Cerri, Irene Fiorilla, Valentina Audrito, Ágata Sofia Assunção Carreira, Alessandro Provenzani

## Abstract

Nicotinamide adenine dinucleotide (NAD⁺) is an essential metabolic cofactor, and maintenance of NAD⁺ homeostasis is critical for cancer cell survival. Pharmacological inhibition of nicotinamide phosphoribosyltransferase (NAMPT), the rate-limiting enzyme of the NAD⁺ salvage pathway, induces NAD⁺ depletion and energetic stress, although the development of resistance limits its therapeutic potential. We have previously shown that acquired resistance to NAMPT inhibition in triple-negative breast cancer (TNBC) cells is associated with extensive metabolic rewiring. Here, we investigated the role of the Hippo pathway effector YAP in the cellular adaptation to NAMPT inhibition. Stable YAP downregulation markedly decreased the sensitivity of MDA-MB-231 cells to the NAMPT inhibitors FK866 and CHS-828, whereas re-expression of constitutively active YAP5SA partially restored sensitivity to FK866. YAP-silenced cells maintained NAD(H) and ATP levels during FK866 exposure and showed reduced oxidative stress and caspase 3/7 activation. Comparative proteomic analysis of YAP-silenced cells and cells with acquired FK866 resistance revealed common alterations in mitochondrial, cell-cycle, and RNA-processing programs. Consistently, YAP downregulation altered the cell-cycle response to FK866 and increased mitochondrial respiration, spare respiratory capacity, and mitochondrial mass. These findings identify YAP as a modulator of the cellular response to NAMPT inhibition and suggest that sustained YAP downregulation contributes to the establishment of an adaptive state characterized by reduced proliferative and biosynthetic programs together with increased mitochondrial capacity.

## INTRODUCTION

Nicotinamide adenine dinucleotide (NAD⁺) is an essential metabolite that couples cellular bioenergetics to signaling and stress responses. It acts as a redox cofactor in glycolysis, the tricarboxylic acid (TCA) cycle and oxidative phosphorylation, while also serving as a substrate for enzymes such as poly(ADP-ribose) polymerases (PARPs), sirtuins and NADases, which regulate DNA repair, transcription and cellular stress responses^1,2^. Maintaining NAD^+^ homeostasis therefore requires continuous biosynthesis through three major routes: the de novo pathway from tryptophan, and two salvage pathways involving nicotinamide phosphoribosyltransferase (NAMPT) and nicotinic acid phosphoribosyltransferase (NAPRT), which use nicotinamide (NAM) and nicotinic acid (NA), respectively, as precursors. Among these, NAMPT represents the rate-limiting enzyme and the major source of NAD⁺ production in mammals^3,4^.

Cancer cells often require increased NAD⁺ availability to sustain their elevated metabolic and biosynthetic demands, maintain redox balance and counteract oxidative stress^5^. Accordingly, NAMPT upregulation has been reported in several solid malignancies and associated with tumor progression, metabolic adaptation, DNA repair, invasion and therapeutic resistance^6,7^. Consequently, NAMPT has emerged as an attractive therapeutic target, with FK866 (APO866) widely investigated in preclinical cancer models. By inhibiting NAMPT, FK866 blocks NAD⁺ biosynthesis, leading to intracellular NAD⁺ and ATP depletion and ultimately cell death^8^. However, prolonged exposure can induce acquired resistance through different mechanisms^9,10^. We have previously showed that triple-negative breast cancer (TNBC) cells acquire resistance to the NAMPT inhibitor FK866 following long-term exposure to the small molecule, which results in a deep reshape of cellular metabolism^11,12^. FK866-resistant cells (RES) exhibit increased glycolytic metabolism, relying on increased lactate dehydrogenase (LDHA) activity, compared with their parental (PAR) counterparts^11^. Further characterization of the model, revealed substantial mitochondrial remodeling in RES cells, including increased mitochondrial mass, maximal respiratory capacity and spare respiratory capacity (SRC), together with enhanced utilization of pyruvate and succinate^12^. However, the signalling mechanisms contributing to the establishment of this adaptive state remain poorly understood.

Yes-associated protein (YAP) and its homologous transcriptional coactivator with PDZ-binding motif (TAZ), the main downstream effectors of the Hippo pathway, are interesting candidates for the regulation of metabolic adaptation. In breast cancer cells, YAP1–TEAD1 promotes glycolysis through the transcriptional regulation of GLUT1^13^, while TAZ/YAP activation enhances glutamine utilization through the induction of GOT1 and PSAT1^14^. In TNBC models, YAP also contributes to mitochondrial redox homeostasis by regulating antioxidant genes and limiting fatty acid oxidation-associated oxidative stress^15^. Importantly, several metabolic processes regulated by YAP/TAZ depend on NAD(H)-dependent enzymatic reactions, suggesting a possible relationship between YAP signaling and cellular adaptation to NAD(H) depletion^16^.

Notably, our previous work in prostate cancer identified a functional relationship between the YAP-nicotinamide N-methyltransferase (NNMT) axis and sensitivity to NAMPT inhibition in stem-like/mesenchymal prostate cancer cells. In this model, YAP or NNMT silencing rescued cells from FK866-induced NAD(H) depletion, cellular stress and apoptosis, highlighting a potential role for YAP in regulating the response to NAD⁺ depletion^17^.

Nevertheless, the contribution of YAP to the metabolic rewiring associated with resistance to NAMPT inhibition in breast cancer remains unexplored. Building on previous characterization of metabolic and mitochondrial remodeling in FK866-resistant MDA-MB-231 cells, here we investigate whether YAP contributes to the establishment of this resistant state and to the cellular adaptations that allow TNBC cells to tolerate NAD⁺ depletion.

## MATERIAL AND METHODS

### Cell lines culture conditions

MDA-MB-231 (MDA) and HEK293T were obtained from American Type Culture Collection (ATCC). MDA shSCR, MDA shYAP, MDA shYAP_empty and MDA shYAP_YAP5SA were obtained as described below. MDA-MB-231 FK866-resistant cell line (RES) was developed by increasing exposure of MDA-MB-231 parental cells (PAR) to FK866 over a 3-month period of time, as previously described^12^. MDA and HEK293T cells were cultured in DMEM (Gibco), supplemented with 10% fetal bovine serum (FBS), 2 mM L-glutamine, and 100 U/mL penicillin-streptomycin (all from Lonza). The culture media of MDA shSCR and MDA shYAP was supplemented with 1 µg/mL of puromycin, while to the one of MDA shYAP_empty and MDA shYAP_YAP5SA was added 500 µg/mL of hygromycin. The media of RES cells were additionally supplemented with 100 nM of FK866.

Human pancreatic carcinoma cell line PK9 was kindly provided by G. Feldmann. PK9 shSCR and PK9 shYAP were obtained as described in ^18^. Cells were grown in RPMI (Gibco) supplemented with 10% FBS, 2 mM L-glutamine, and 100 U/mL penicillin-streptomycin (all from Lonza). The culture media of PK9 shSCR and PK9 shYAP was supplemented with 1 µg/mL of puromycin.

All cell lines were kept under humidified conditions with 5% CO_2_ at 37 °C.

### Drug treatment and in vitro cell viability

Cell lines were treated with FK866 (sc-205325, Santa Cruz Biotechnology) or CHS-828 (200484-11-3, Cayman Chemical), for the indicated times. For *in vitro* drug sensitivity assessment, cells were treated for 48 hours with FK866. The OzBlue Cell Viability kit (OzBiosciences) was used to measure the compound toxicity after 2 hours of incubation, according to the manufacturer instructions. Sulforhodamine B (SRB) assay was carried afterwards in the same plate. Cells were fixed with 10% trichloroacetic acid (TCA) for 30 minutes at 4 °C, and after which washed with cold Milli-Q water. Cells were stained with SRB (0.4% SRB (S1402, Merck) in 1% acetic acid) for 10 minutes and washed with 1% acetic acid solution. After drying, the stain was extracted with 10 mM Tris Base and absorbance was measured at 560 nM (Abs_560_). The Abs_560_ is proportional to the number of cells in each condition.

### Caspase 3/7 activation assay

MDA shSCR and MDA shYAP were seeded into tissue cultured transparent clear-bottom 96-well plates and allowed to adhere for 24 hours. Cells were then treated with the indicated concentrations of FK866 and CellEvent^TM^ Caspase 3/7 Green (Life Technologies, C10432) for 72 hours, as described in ^19^. Data was collected with IncuCyte® S3 Live-Cell Analysis System (Sartorius) in both brightfield and FITC channels. Images were analyzed for number of green objects per well. Results are presented as a fold change of the respective non-treated conditions.

### Reactive Oxygen species (ROS) levels determination

Cellular ROS production was detected by flow cytometry (Sysmex-partec) using 5-(and 6-)-chloromethyl-2′,7′-dichlorodihydrofluorescein diacetate (CM-H_2_DCFDA, Thermo Fisher Scientific), a chloromethyl derivative of H_2_DCFDA, useful as an indicator for ROS in cells. Briefly, cells were resuspended in pre-warmed PBS, loaded with 1 μM CM-H_2_DCFDA 5-(and 6-)-chloromethyl-2′,7′-dichlorodihydrofluorescein diacetate, incubated in the dark for 30 minutes at 37 °C, 5% CO_2_. Then cells were washed with PBS and incubated for 15 minutes. The oxidative conversion of 5-(and 6-)-chloromethyl-2′,7′-dichlorodihydrofluorescein diacetate to its fluorescent product was measured immediately by flow cytometry. Data are reported as percentage of ROS positive cells.

### Colony formation assay (soft agarose)

Soft agar assay was carried out in 6-well plates with a double coating of agarose. The base agarose layer presented 0.7% of agarose in 10% FBS supplemented DMEM. For the top layer, 5000 cells per condition were resuspended in 1 mL of 0.35% agarose in 10% FBS supplemented DMEM and seeded. Cells were covered with 1 mL of complete media and treated with different concentrations of FK866 every 3 days from day 2, for 15 days; and constantly kept under selection with puromycin (1 µg/mL). Total colony number and colony diameter were measured using Operetta and Harmony 3.5.2 software.

### Determination of NAD(H) and ATP levels

Both NAD(H) and ATP levels were assessed following 48 hours of FK866 exposure. Intracellular NAD(H) was measured with the NAD/NADH-Glo (G9071, Promega), while ATP levels were measured with the CellTiter-Glo Luminescent Cell Viability Assay (G7571, Promega), according to the manufacturer’s protocol. The SRB assay was performed under the same conditions and used for normalization of the luminescence values.

### Protein extraction and western blot analysis

Samples were lysed for 10 min on ice with RIPA lysis buffer supplemented with Protease Inhibitor Cocktail (ThermoScientific) and quantified with the Pierce™ BCA Protein Assay (23225). The SDS-polyacrylamide gel was loaded with equal amounts of proteins (20 µg) and used for protein separation, followed by the transference to a PVDF membrane. The primary antibodies used were: YAP (Santa Cruz Biotechnologies. sc-101199), p-YAP (Cell Signaling Technologies, 4911S) and β-actin (BK4970S CST). β-actin was used as protein loading controls. The anti-mouse and anti-rabbit secondary antibodies (111-035-003) used were obtained from Jackson Immunoresearch Laboratories. Signal was revealed with chemiluminescence detection kit reagents (Amersham ECL Select, GE Healthcare) at ChemiDoc (BioRad). A representative western blot of three independent biological experiments is shown.

### RNA extraction and real-time qPCR

Total RNA extraction was performed with TriZOL reagent (Invitrogen), followed by cDNA synthesis (RevertAid RT Reverse Transcription Kit, K1691, ThermoScientific), all according to the manufacturer’s instructions. Real-time quantitative PCR (RT-qPCR) analyses were performed in technical triplicates using the ExcelTaq™ 2X qPCR Master Mix (SMOBIO) on a CFX96 Real-Time PCR Detection System (Bio-Rad). The sequences of the primers used were: PPARGC1A FW – 5’ CCAAAGGATGCGCTCTCGTTCA 3’ and PPARGC1A RV – 5’ RV: CGGTGTCTGTAGTGGCTTGACT 3’. ΔCq method was used for samples’ mRNA quantification with RPLP0 as the housekeeping gene.

### Stable and transient YAP silencing

Lentiviral vector particles (LVP) for shYAP or YAP5SA were produced in HEK293T cells, using pLKO or pQCXIH-Myc vectors as the transfer plasmids, respectively. Cells were co-transfected with a Δ891 and a VSV-G encoding vector along with the shYAP or YAP5SA plasmids. Viral supernatants were collected and filtered 72 h post-transfection. Following the quantification with the PERT assay, 1 Reverse Transcription Unit (RTU) was used to infect MDA cells. Selection with puromycin (1 µg/mL) was performed on MDA cells stably transduced with shYAP or YAP5SA. The respective scramble controls using the same plasmid backbone (either pLKO or pQCXIH-Myc) were also produced.

For YAP transient silencing, PAR cells were transfected with either control siRNA (sc-37007) or YAP siRNA (sc-38637), at a concentration of 5nM. Following 48 hours of silencing, cells were treated for additional 48 hours with FK866 at the indicated concentrations. Lipofectamine 3000 (Invitrogen, L3000015) was used for the transfection reaction, following the manufacturer’s instructions.

### Mitochondrial function

The mitochondrial function (mitochondrial respiration) was assessed in MDA shSCR and MDA shYAP with the Seahorse MitoStress kit (103015-100, Agilent), following the manufacturer’s instructions. Shortly, cells were seeded in triplicate for 48 hours in seahorse assay plates in Seahorse DMEM supplemented with 10 mM glucose, 2 mM glutamine, 1 mM pyruvate (all from Agilent). The day before the assay, a cartridge was hydrated using Seahorse calibrant XF (Agilent) and incubated overnight at 37 °C in a CO_2_ deprived atmosphere. The oxygen consumption rate (OCR) was determined on a XFe96 Seahorse Bioanalyzer (Agilent). The final concentration of injected compounds was: 1 µM oligomycin; 2 µM FCCP (carbonyl cyanide-4(trifluoromethoxy) phenylhydrazone) and 5 µM of each rotenone and antimycin A. OCR was normalized to the samples’ protein content using the Bradford reagent.

### Mitochondrial mass determination

The MitoTracker Green FM (Life Technologies, M7514) was used to determine the mitochondrial mass of MDA shSCR and shYAP. 200 cells per well were seeded in 384-well optic glass plates (CellCarrier384, Perkin Elmer), and incubated with MitoTracker Green FM (100 nM) and Hoechst (1:5000) for mitochondria and nuclei staining, respectively. Z-stacks were acquired with a Leica TCS SP8 laser scanning confocal system, installed on a Leica DMi8 inverted fluorescence microscope, using a HC PL APO 20x/0.75 CS2 objective. The acquisition parameters were kept constant throughout all the conditions.

The mean intensity of the green fluorescent MitoTracker signal was quantified after single-cell segmentation, on the sum projection of all the stacks comprising the total area of each cell. Both cell segmentation and signal intensity quantification were obtained from a semi-automatic macro in Fiji/ImageJ. Representative images were linearly adjusted only for brightness and contrast. At least 300 nuclei were analyzed per condition.

### Sample preparation for LC-MS analysis

MDA shSCR, MDA shYAP, PAR and RES cells and the respective FK866 treated conditions were suspended in 50 mM HEPES pH 8.0, 2% SDS, supplemented protease inhibitor and Benzonase (500 U/mL), and incubated for 30 min at 4 °C under rotation. After that, samples were measured for protein content using the Pierce™ BCA Protein Assay (23225) assay. For each sample, proteins (30 µg) were reduced with 10mM DTT at 56 °C for 30 minutes. Alkylation of cysteine residues was achieved by adding 22.5 mM iodoacetamide (IAA) for 30 min in the dark at room temperature. The reaction was then quenched with 10 mM DTT.

Protein clean-up, digestion, and peptide clean-up were performed using Single-pot solid-phase-enhanced sample preparation (SP3) protocol, as previously described^20^. 6 μL of washed magnetic beads (1:1 hydrophilic and hydrophobic beads at 50 µg/µL) were added to the lysates. Afterward, acetonitrile (ACN) was added to a final concentration of 70% (v/v), and samples were mixed at room temperature for 18 min. Beads were then immobilized by incubation on a magnetic rack for 2 min. The supernatant was discarded, and the beads were washed twice with 70% (v/v) EtOH and once with 100% ACN. The beads were suspended in 45 μL of digestion buffer (100 mM NH_4_HCO_3_, 5 mM CaCl_2_ supplemented with 1.2 µg of Trypsin), and incubated overnight at 37 °C on a mixing platform. After the overnight digestion, the samples were collected by incubation on a magnetic rack for 2 min and subjected to peptide clean-up. Acetonitrile (100%) was added to each tube to achieve a final concentration > 95%. The mixtures were mixed for 5 min at room temperature and placed on a magnetic rack for 2 min. The supernatant was discarded, and the beads were rinsed once with 500 μL of 100% ACN. The beads were then suspended in 2% DMSO, sonicated for 1 min, and mixed on a shaker for 5 min. Peptides were collected by immobilizing the beads on a magnetic rack for 2 min. The supernatant containing purified peptides was transferred into a fresh tube. Before LC–MS analysis, the peptides were acidified using 0.1% FA.

### LC-MS analysis

Digested samples were separated using an Easy-nLC 1200 system (Thermo Scientific). A 30 cm reversed-phase C18 column (inner diameter 75 µm, 1.7µm particle size, MSWIL, the Netherlands) heated at 40 °C was employed for peptide separation, with a two-component mobile phase system consisting of 0.1% formic acid in water (buffer A) and 0.1% formic acid in 80% acetonitrile (buffer B). Peptides were eluted using a gradient of 5% to 25% over 55 minutes, followed by 25% to 40% over 10 minutes, by 40% to 98% over 10 minutes, and finally kept at 98% over 10 minutes. The flow rate was set to 200 nL/min.

Samples were injected into an Orbitrap Fusion Tribrid mass spectrometer (Thermo Scientific, San Jose, CA, USA) and data were acquired in data-dependent mode (2100 V). The temperature of the ion transfer tube was maintained at 200 °C. Full scans were performed at a resolving power of 120.000 FWHM (at 200 m/z) with an AGC target of 1x10e6. The mass range of 350-1100 m/z was surveyed for precursors, with first mass set at 140 m/z for fragments. Each full scan was followed by a set of MS/MS scans (HCD, collision energy of 30%) with a 3-second cycle time and a maximum injection time of 150 ms in the ion trap, along with an AGC target of 5x10e3. A dynamic exclusion filter was set to apply every 40 seconds with a ± 5 ppm mass tolerance. Data were acquired using Thermo software Xcalibur (version 4.3) and Tune (version 3.3). During the project, QCloud^21^ was used to monitor instrument longitudinal performance.

Peptides searches were conducted in Proteome Discoverer 2.2 software (Thermo Fisher Scientific) against the Homo sapiens FASTA file (Uniprot, downloaded in March 2023) and a database containing common contaminants. Proteins were identified using MASCOT search engine with a mass tolerance of 10 ppm for precursors and 0.6 Da for products. Trypsin was selected as the enzyme with a maximum of 5 missed cleavages. Static modification of carbamidomethyl (C) and variable modification of oxidation (M) and acetylation (protein N-term) were incorporated in the search. False discovery rate filtering was applied, with a threshold set to < 0.05 at the PSM, peptide and protein levels. Results were filtered to exclude potential contaminants and proteins with fewer than two peptides.

### MS computational analyses

MS downstream analysis was conducted using the ProTN proteomics pipeline (www.github.com/TebaldiLab/ProTN and www.rdds.it/ProTN). In summary, peptide intensities were log2-transformed, normalized using median normalization and summarized into proteins through the utilization of functions in the DEqMS Bioconductor package^22^. Imputation of the missing intensities was performed using PhosR package^23^. Differential analysis was carried out with the DEqMS package, where proteins with an absolute log2 fold change (FC) greater than 0.75 and p-value less than 0.05 were deemed significant. Functional enrichment analysis of differentially expressed proteins was accomplished using ClusterProfiler function EnrichGO^24^. Enriched terms with p-value lower than 0.05 were considered significant.

### MS data availability

The mass spectrometry proteomics data have been deposited to the ProteomeXchange Consortium via the PRIDE^25^ partner repository with the dataset identifier PXD046817 and 10.6019/PXD046817.

### Cell cycle

The DNA staining propidium iodide (PI) was used for DNA content-based assessment of cell cycle distribution. Cells were treated for 48 hours with FK866, fixed with 70% ethanol and stained with 10 µg/mL of PI. RNAse (100 µg/mL) was added to the staining solution to ensure optimal DNA resolution, considering that PI can also bind RNA. Flow cytometry was conducted on the BD FACSymphony^TM^ A1 and acquired by the BD FACSDiva^TM^ Software. Starvation for 24 hours and 2 hours treatment with Nocodazole (1 µg/mL) were used as positive controls, respectively for G1 and G2/M blockage. The percentage of cells in G1, S and G2/M phases was calculated with the software ModFit LT 4.1.

### Statistical Analysis

Collected data was expressed as mean ± standard error of mean (SEM) of three independent biological experiments conducted in technical triplicates, unless indicated otherwise. Statistical significance between control and test groups was determined using Student’s t-test or one-way ANOVA, considering values of *p < 0.05, **p < 0.01, and ***p < 0.001 as significant. Data were plotted in Graph prism 6 software.

## RESULTS

### Sustained YAP downregulation confers resistance to NAMPT inhibition

Our previous studies characterized the metabolic adaptations sustaining the acquired resistance to NAMPT inhibition in TNBC cells^11,12^. More recently, we have demonstrated that YAP activity determines the innate sensitivity of mesenchymal stem-like castration-resistant prostate cancer cells to NAMPT inhibition^17^. We therefore hypothesized that sustained reduction of YAP activity could promote a cellular state with decreased sensitivity to NAMPT inhibition in TNBC.

MDA-MB-231 cells were stably silenced for YAP (MDA shYAP) and exposed to the NAD(H) depletion agent FK866, a selective inhibitor of NAMPT. YAP reduced expression in MDA shYAP was confirmed by western blot (Figure 1A). Cellular viability was assessed upon 48 hours of treatment, with the OzBlue assay, and revealed a decrease of sensitivity to FK866 in cells silenced for YAP, compared with the scramble condition (IC_50_ = 6 nM) (Figure 1B). This observation was validated in an independent cellular model of pancreatic ductal adenocarcinoma (PK9), in which it was confirmed a decreased sensitivity to FK866 in cells stable silenced for YAP (PK9 shYAP – IC_50_ = 605,4 nM) compared with the scramble condition (PK9 shSCR - IC_50_ = 52,30 nM) (Supplementary Figure 1A-B).

**Figure 1:**
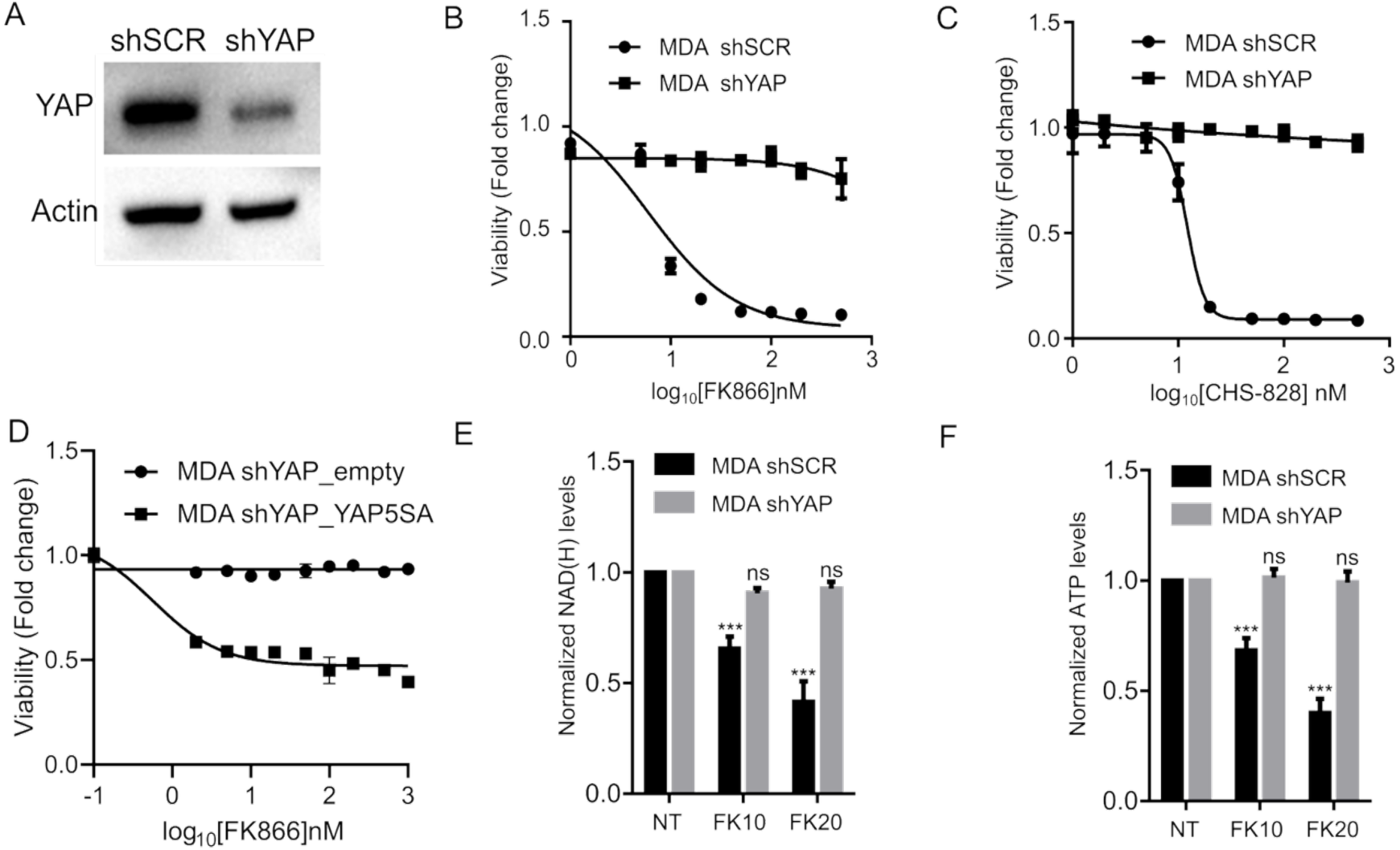
YAP silencing renders MDA cells resistant to NAMPT inhibition. (A) The silencing of YAP in MDA cells was confirmed by western blotting, with actin used as a loading control. The viability of MDA shSCR and MDA shYAP upon 48 hours of exposure to the NAMPT inhibitors FK866 (B) and CHS-828 (C), with ranging concentrations from 1 to 500 nM, was assessed by the OzBlue viability assay. (D) Viability of MDA shYAP_empty and MDA shYAP_YAP5SA upon 48 hours of exposure to the FK866, assessed by the OzBlue assay. (E) NAD(H) and (F) ATP levels were quantified in MDA shSCR and MDA shYAP, exposed to 10 or 20 nM of FK866 for 48 hours. The results were normalized to the protein content quantified with the SRB assay. All the experiments were performed in triplicate. Repeated measures one-way ANOVA was used to calculate statistical significance (ns – not significant; *** *P*<0.001) between non treated (NT) and experimental conditions.

To understand whether the observed FK866 acquired resistance was specific, the sensitivity to a second NAMPT inhibitor (CHS-828) was also tested. The IC_50_ value of 12.2 nM for MDA shSCR was obtained upon 48 hours of treatment (Figure 1C). To exclude the possibility that this phenotype was influenced by the use of a metabolism-dependent viability assay, drug sensitivity was independently evaluated using the sulforhodamine B (SRB) assay. Comparable responses were observed, with IC_50_ values of 10 nM and 13 nM for MDA shSCR, respectively after treatment with FK866 and CHS-828 (Supplementary Figure 1C-D). Considering the lack of sensitivity of MDA shYAP to NAMPT inhibition, it was not possible to calculate its IC_50_ with any of the assays (Figure 1C-D; Supplementary Figure 1C-D). Interestingly, the transient silencing of YAP by RNA interference only partially reduced FK866 sensitivity in PAR cells, suggesting that sustained YAP downregulation may be required for cells to develop adaptative responses to increase the tolerance to NAD depletion (Supplementary Figures 1E-F).

To establish whether the resistant phenotype induced by stable YAP downregulation was specifically attributable to loss of YAP activity, we performed a rescue experiment by expressing the constitutively active YAP mutant YAP5SA in MDA shYAP cells. YAP5SA carries substitutions at five regulatory phosphorylation sites, preventing inhibitory phosphorylation by upstream kinases and promoting constitutive nuclear YAP activity. MDA shYAP overexpressing YAP5SA (MDA shYAP_YAP5SA) restored sensitivity to FK866 (Fig 1D), compared with the corresponding empty-vector control. These results demonstrate that resistance to FK866 following stable YAP downregulation is YAP-dependent and can be reversed by restoration of YAP activity.

We next asked whether the differential response to FK866 was reflected in the ability of cells to maintain their energetic state during NAMPT inhibition. Depletion of NAD(H) and ATP levels in a dose-dependent fashion was observed upon 48 hours of exposure to the small molecule in MDA shSCR (Figure 1E-F). Differently, no modulation of either NAD(H) or ATP levels were observed in MDA shYAP, confirming the lack of sensitivity to the small molecule (Figure 1E-F). Together, these findings show that sustained YAP downregulation confers resistance to NAMPT inhibition and enables MDA-MB-231 cells to maintain NAD(H) and ATP pools during FK866 exposure. Conversely, restoration of YAP activity re-establishes FK866 sensitivity, identifying YAP as a determinant of the cellular response to NAMPT inhibition.

### YAP silencing prevents FK866-induced apoptosis activation and preserves anchorage independent growth

NAMPT inhibition is described to induce strong increase in cellular reactive oxygen species (ROS), ultimately inducing apoptosis in different cancer models^26,27^. We next therefore investigated whether the FK866-resistance conferred by YAP downregulation was associated with and altered oxidative stress response. Flow cytometry analysis revealed that the percentage of ROS positive MDA shYAP cells is significantly lower than the MDA shSCR ones, suggesting that YAP downregulation is associated with reduced oxidative stress (Figure 2A). To assess whether this phenotype was accompanied by reduced apoptosis, we quantified the activation of caspase 3/7, upon exposure to FK866 for 72 hours by live-cell imaging. Consistent with their sensitivity to NAMPT inhibition, MDA shSCR cells showed a concentration-dependent increase in caspase 3/7 activation, reaching approximately two-fold relative to the untreated condition (Figure 2B).

**Figure 2:**
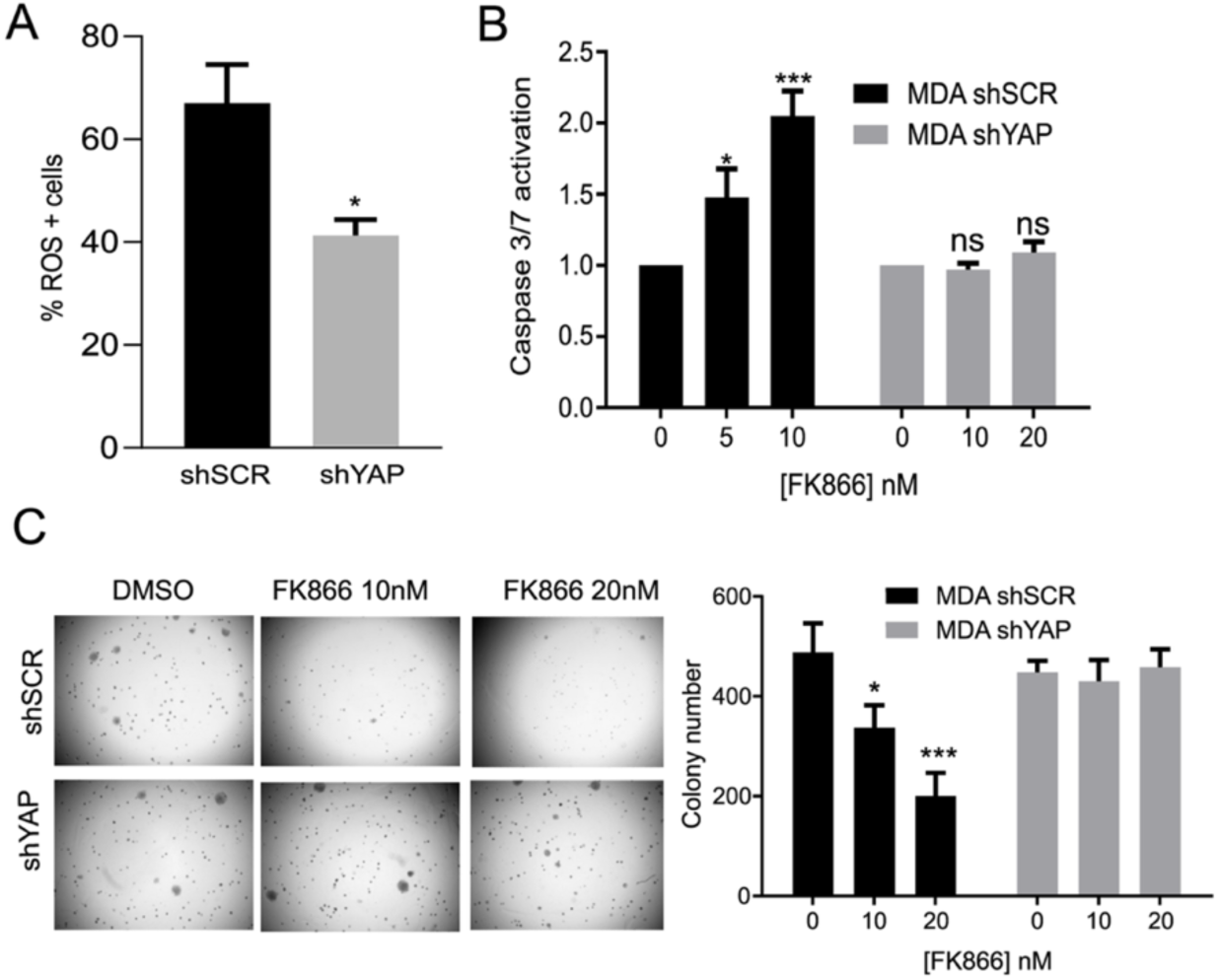
MDA shYAP cells present less ROS positive cells and apoptosis induction upon NAD depletion, and are able to grow in anchorage-independent conditions. (A) Quantification of the percentage of ROS positive cells in MDA shSCR and MDA shYAP, with CM-H2DCFDA dye by flow cytometry. (B) Quantification of caspase 3/7 activation upon 72 hours of FK866 (FK) exposure at the indicated concentrations (unit: nM) in MDA shSCR and MDA shYAP. Results are presented as a fold change to non treated condition for each cell line. (C) Representative images of soft agarose assay of MDA shSCR and MDA shYAP. Cells were treated with different concentrations of FK866 (5 nM, 10 nM, 20 nM) every three days for a period of three weeks. DMSO was used as negative control. Quantification of normal colony number exposed to the described concentrations of FK866. Colony number was quantified with Operetta and analyzed on Harmony 3.5.2 software. Three biological replicates were performed (\**P*< 0.05, *** *P*< 0.001 compared to DMSO condition).

Instead, no significant increase of apoptosis was observed in MDA shYAP, despite exposure to higher FK866 concentration (Figure 2B). Thus, YAP downregulation is associated with reduced ROS levels and impaired activation of the apoptotic response to FK866.

Given the established role of YAP in anchorage-independent growth^28^, we next investigated whether the resistance observed in two-dimensional cultures was maintained under three-dimensional growth conditions. MDA shSCR and MDA shYAP were grown in soft agar for 2 weeks, with drug supplementation every 3 days. FK866 treatment significantly reduced MDA shSCR colony number in a dose-dependent manner (Figure 2C). In contrast, MDA shYAP colonies’ number was independent of FK866 exposure, confirming their lack of sensitivity to the small molecule (Figure 2C). Importantly, no significant difference was observed in the non-treated conditions, meaning that YAP silencing *per se* does not affect colony-forming capacity under these experimental conditions.

Overall, these data reveals YAP as a modulator of FK866 sensitivity in MDA cells, whose genetic downregulation prevents ROS accumulation and apoptosis upon exposure to the small molecule, while maintaining NAD(H) and ATP levels to sustain both in attachment conditions in 2D culture and anchorage-independent cellular growth.

### Proteomic analysis of MDA shYAP and RES reveals cell cycle deregulation and mitochondrial dependency as common FK866-resistance adaptative programs

The stronger resistance observed following stable compared with transient YAP downregulation suggested that sustained YAP loss may promote cellular adaptations that increase tolerance to NAD depletion. We therefore asked whether the resistant state induced by YAP downregulation recapitulates molecular features of the acquired FK866 resistance previously characterized in MDA-MB-231 cells^12^. To address this, we performed label-free quantitative proteomic analysis of MDA shYAP cells relative to MDA shSCR controls and compared the resulting proteomic changes with those observed in FK866-resistant (RES) cells relative to their parental FK866-sensitive counterpart (PAR).

A total of 4315 proteins were identified, of which 823 were significantly differentially expressed in MDA shYAP compared with the MDA shSCR. Among these, 451 proteins were upregulated and 372 were downregulated (Figure 3A; Supplementary Material 1). As expected, YAP abundance was reduced in MDA shYAP cells (log_2_FC = −0.71; p = 0.008). Interestingly, NAMPT, the direct target of FK866, was also significantly downregulated (log_2_FC = −1.21; p = 3.5 × 10^−8^) (Figure 3A). Gene ontology analysis (GOA) of proteins upregulated following stable YAP silencing revealed enrichment of mitochondrial and oxidative-metabolism-related processes, such as “energy derivation by oxidation of organic compounds”, “cellular respiration” and “aerobic respiration”, consistent with a shift towards increased mitochondrial metabolism in MDA shYAP cells (Figure 3B). In contrast, proteins downregulated following YAP silencing were enriched for cell cycle and mitotic processes, including “chromosome segregation” and “mitotic nuclear division” (Figure 3C), suggesting altered regulation of cell-cycle progression. Processes related to RNA metabolism and protein synthesis, including “ribonucleoprotein complex biogenesis”, “mRNA processing” and “RNA splicing”, were also enriched among downregulated proteins (Figure 3C), consistent with our previous observations of altered translational control in response to NAMPT inhibition^29^.

**Figure 3:**
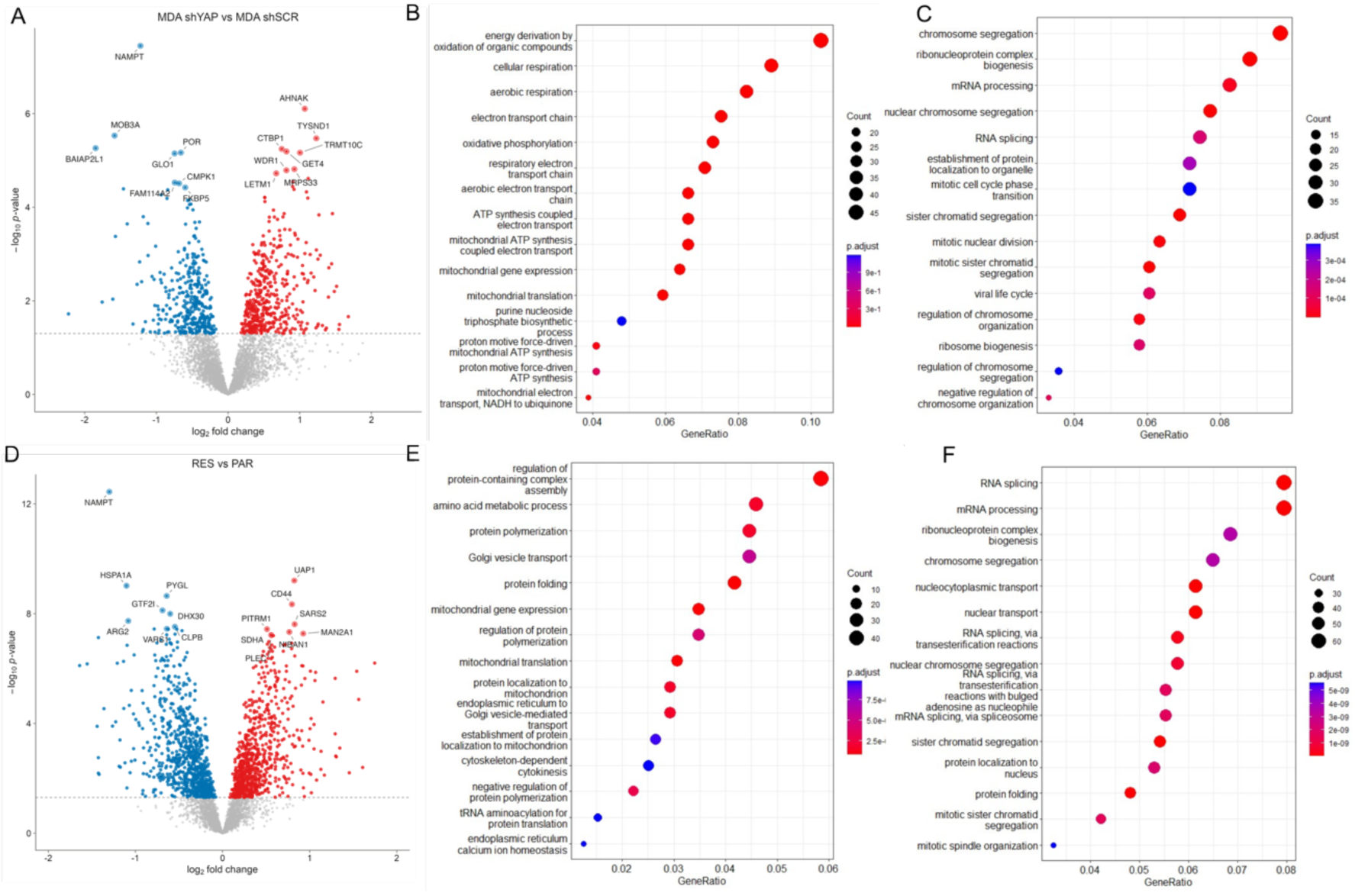
Proteomic analysis reveals common mitochondrial and cell-cycle alterations in MDA shYAP and RES cells. (A) Volcano plot showing differentially expressed proteins in MDA shYAP compared with MDA shSCR cells. Gene Ontology analysis (GOA) of proteins significantly upregulated (B) and downregulated (C) in MDA shYAP cells. (D) Volcano plot showing differentially expressed proteins in RES compared with PAR cells. GOA of proteins significantly upregulated (E) and downregulated (F) in RES cells. The top 15 enriched biological processes are shown for each comparison. In the volcano plots, significantly upregulated and downregulated proteins are shown in red and blue, respectively, while non-significantly altered proteins are shown in grey. In the GOA plots, GeneRatio indicates the proportion of differentially expressed proteins associated with each GO term, dot size represents the number of differentially expressed proteins associated with the respective term, and color indicates the adjusted p-value.

We next asked whether cells that had acquired resistance through prolonged FK866 exposure displayed a similar proteomic profile. A total of 4316 proteins were identified, of which 1644 were significantly differentially expressed in RES cells compared with PAR controls, including 779 upregulated and 865 downregulated proteins (Figure 3D; Supplementary Material 1). Notably, NAMPT was also strongly downregulated in RES cells (log_2_FC = −1.30; p = 4.16 × 10^−1^^3^) (Figure 3D), identifying reduced NAMPT abundance as an additional feature shared between of resistance associated with sustained YAP downregulation and resistance acquired through chronic FK866 exposure.

GOA of proteins upregulated in RES cells revealed a prominent mitochondrial signature, including processes associated with “mitochondrial gene expression” and “mitochondrial translation” (Figure 3E). Among the upregulated proteins was succinate dehydrogenase assembly factor 2 (SDHAF2, log_2_FC = 1.74; p = 6.3 × 10^−7^), consistent with our previous demonstration that RES cells develop increased dependence on succinate metabolism^12^. In contrast, proteins downregulated in RES cells were enriched for cell-cycle and mitotic processes, including “chromosome segregation”, “mitotic sister chromatid segregation” and “mitotic spindle organization” (Figure 3F). RNA-processing programs, including “RNA splicing”, “mRNA processing” and “ribonucleoprotein complex biogenesis”, were also enriched among downregulated proteins (Figure 3F).

Overall, proteomics data point to a pathway-level convergence in both cellular models of resistance to NAMPT inhibition, sustained by cell cycle and translation deregulation, as well as an increased mitochondrial metabolism dependency.

### YAP downregulation and acquired FK866 resistance alter the cell-cycle response to NAD depletion

Given the consistent downregulation of cell-cycle and mitotic programs identified by the proteomic analysis in both MDA shYAP and RES cells, we next investigated whether these changes were associated with a shared alteration in the cell-cycle response to NAMPT inhibition. Cell-cycle distribution was assessed by propidium iodide (PI) staining and flow cytometry, allowing quantification of cells in G0/G1, S and G2/M phases based on DNA content. Following 48 hours of FK866 exposure, MDA shSCR cells showed a concentration-dependent alteration in cell-cycle distribution, with a pronounced accumulation in G2/M at 10 nM FK866. The proportion of MDA shSCR cells in G2/M increased from 15.2% under untreated conditions to 31.1% following treatment with 10 nM FK866, accompanied by a decrease in the G0/G1 population from 49.1% to 35.0% (Figure 4A). In contrast, FK866 exposure did not significantly alter the distribution of MDA shYAP cells across G0/G1, S and G2/M phases, even at 100 nM FK866 (Figure 4A).

**Figure 4:**
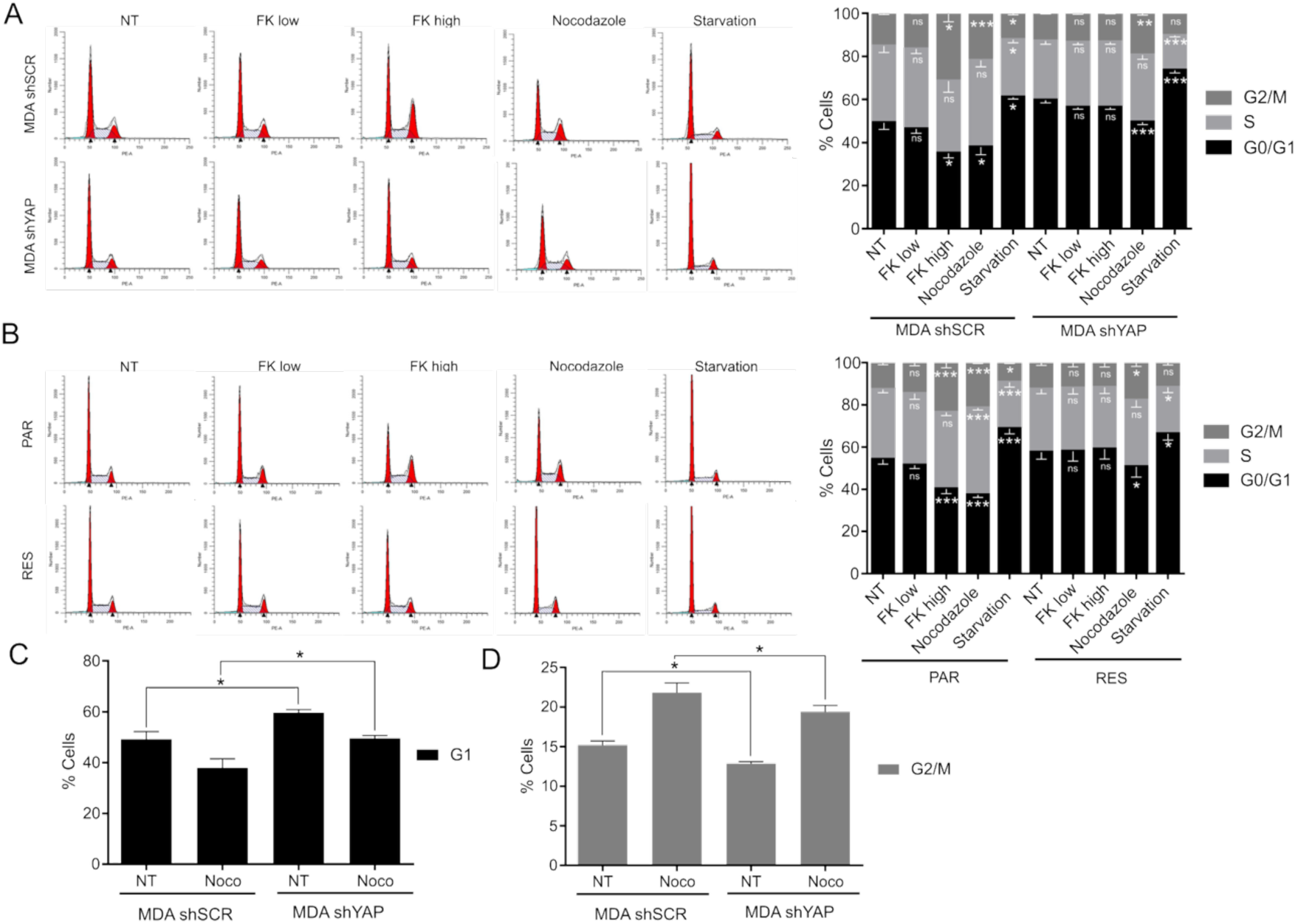
MDA shYAP and RES present deregulated cell cycle checkpoints. (A) Representative images and quantification of MDA shSCR and shYAP (A) and PAR and RES (B) cell cycle distribution, analyzed with the ModFit software. Quantification of the effect of Nocodazole treatment in the G1 (C) and G2/M (D) phases of the cell cycle in MDA shSCR/shYAP cells. For all the experiments, cells were treated for 48 hours with low or high concentrations of FK866, before fixation and PI staining (low: 5 nM for MDA shSCR/PAR and 10 nM for MDA shYAP/RES; high: 10 nM for MDA shSCR/PAR and 100 nM for MDA shYAP/RES). Treatment of Nocodazole (1 µM) for 2 hours was used as a positive control for cell cycle arrestment in G2/M phases. Overnight starvation was used as a positive control for cell cycle arrestment in G1 phase. Three biological replicates were performed for cell cycle analysis of MDA shSCR and MDA shYAP. Five biological replicates were performed for cell cycle analysis of PAR and RES (ns – not significant, * *P*< 0.05, ** *P* < 0.01, *** *P* < 0.001 compared to non treated – NT - condition).

A similar differential response was observed between PAR and RES cells. In PAR cells, treatment with 10 nM FK866 increased the proportion of cells in G2/M from 12.7% to 23.4%, while the G0/G1 population decreased from 54.1% to 40.2% (Figure 4B). In contrast, RES cells did not show FK866-induced accumulation in G2/M, even when exposed to 100 nM FK866 (Figure 4B). Thus, both stable YAP downregulation and acquired FK866 resistance are associated with loss of the G2/M accumulation observed following NAMPT inhibition in sensitive cells.

Cells were additionally subjected to serum starvation or nocodazole treatment as controls for G0/G1 and G2/M accumulation, respectively (Figure 4A-B). Serum starvation increased the proportion of cells in G0/G1, whereas nocodazole increased the G2/M population across the cell models (Figure 4A-B). However, the magnitude of the response to nocodazole differed in MDA shYAP cells, which showed a significantly lower proportion of cells in G2/M and a correspondingly higher proportion in G0/G1 (Figure 4C-D). Interestingly, stable YAP downregulation also altered cell-cycle distribution under basal conditions. Untreated MDA shYAP cells displayed a significantly higher proportion of cells in G0/G1 and a lower proportion in G2/M compared with MDA shSCR cells (Figure 4C-D). Together with the attenuated response to nocodazole, these findings indicate that sustained YAP downregulation alters cell-cycle regulation independently of FK866 exposure.

### YAP silencing increases mitochondrial function and mass

Our previous work identified mitochondrial rewiring as a key feature of acquired resistance to FK866 in MDA-MB-231 cells^13^. Given the enrichment of mitochondrial and oxidative metabolism-related programs identified by proteomic analysis following stable YAP downregulation (Figure 3B), we next investigated whether this signature was accompanied by functional changes in mitochondrial respiration.

Mitochondrial function was assessed by real-time monitoring of the oxygen consumption rate (OCR) following sequential injection of oligomycin, FCCP, and rotenone/antimycin A (Figure 5A). Oligomycin inhibits ATP synthase (complex V), whereas FCCP uncouples oxidative phosphorylation, allowing determination of maximal respiratory capacity. The difference between maximal and basal respiration defines the spare respiratory capacity, reflecting the ability of cells to increase mitochondrial respiration in response to increased energetic demand. MDA shYAP cells displayed higher basal and maximal respiration compared with MDA shSCR cells, together with increased spare respiratory capacity (Figure 5A), demonstrating that sustained YAP downregulation is associated with enhanced mitochondrial respiratory function.

**Figure 5:**
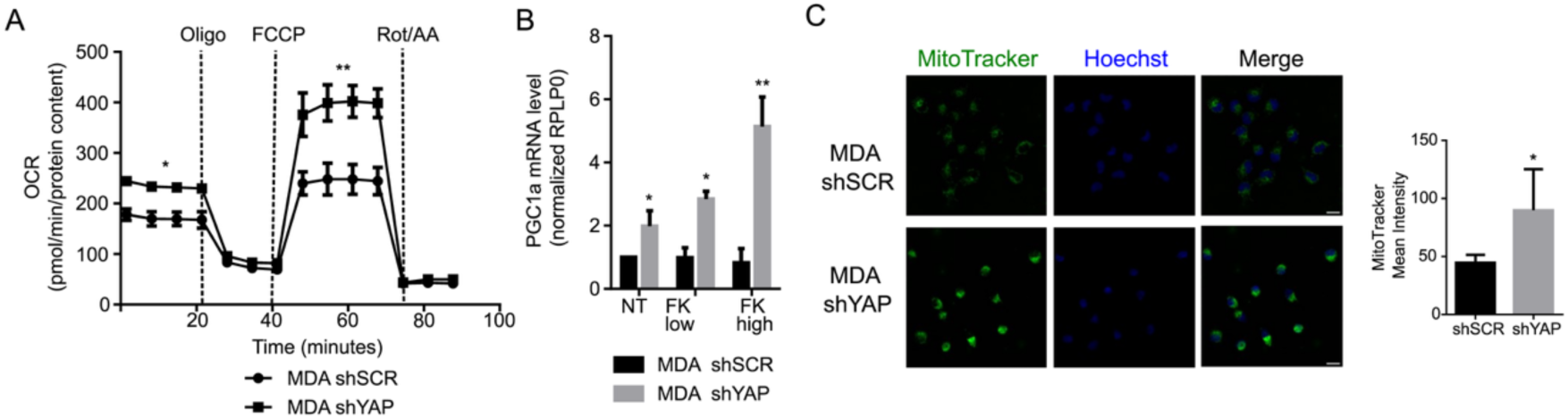
YAP silencing increases mitochondrial function and mass. (A) Representative MitoStress test to determine the oxygen consumption rate (OCR) of MDA shSCR and MDA shYAP upon mitochondrial function modulators injection over time. Abbreviations: Oligo - Oligomycin; FCCP-Carbonyl cyanide-p-trifluoromethoxyphenylhydrazone; Rot/AA - Rotenone and Antimycin A. (B) RT-qPCR to determine PGC1-α expression in MDA shSCR and MDA shYAP, as a proxy of mitochondrial biogenesis. Cells were treated with low or high concentrations of FK866 for 48 hours (low: 5 nM for MDA shSCR and 10 nM for MDA shYAP; high: 10 nM for MDA shSCR and 100 nM for MDA shYAP). *RPLP0* was used as a house-keeping gene. (C) Mitotracker Green FM staining of MDA shSCR and MDA shYAP cells’ mitochondria (Green - left) and Hoechst (Blue-middle) staining of cells nuclei. Composite image as a merge of the green and blue channels with scaling bar of 20 µm (right). Quantification of the MitoTracker mean intensity of at least 300 nuclei of MDA shSCR and MDA shYAP cells is shown. All the experiments were performed in triplicate. Repeated measures one-way ANOVA was used to calculate statistical significance (ns-not significant, \**P*<0.05, \*\**P*<0.01) between MDA shSCR and MDA shYAP.

We next investigated whether the increased respiratory capacity of MDA shYAP cells was accompanied by changes in mitochondrial mass, as previously observed in RES cells. Expression of *PPARGC1A*, encoding the transcriptional coactivator PGC-1α, a major regulator of mitochondrial biogenesis, was significantly increased in MDA shYAP cells compared with MDA shSCR controls (Figure 5B). Moreover, *PPARGC1A* expression increased further following FK866 exposure in MDA shYAP cells (Figure 5B). To directly assess mitochondrial mass, cells were stained with MitoTracker Green, which accumulates in mitochondria largely independently of mitochondrial membrane potential, together with Hoechst for nuclear identification. Quantitative image analysis revealed significantly higher MitoTracker Green fluorescence intensity in MDA shYAP cells compared with MDA shSCR controls (Figure 5C), consistent with increased mitochondrial mass. Together with the increased respiratory capacity, these findings show that sustained YAP downregulation recapitulates key features of the mitochondrial phenotype previously associated with acquired FK866 resistance.

## DISCUSSION

Metabolic reprogramming is a general term for the multiple, coordinated pathways that converge in cellular adaptation to energy deprivation. NAD^+^ is an essential cofactor for both cytoplasmic and mitochondrial metabolic reactions, and consequently metabolic adaptation to NAD^+^ deprivation is critical for cell survival upon NAMPT inhibition^30^.

We have previously demonstrated that cancer cells can acquire resistance to NAMPT inhibition through extensive metabolic rewiring, including increased mitochondrial fitness and altered utilization of metabolic substrates^11,12^. However, the signalling mechanisms contributing to this adaptation are not fully understood.

Here, we showed that YAP is a modulator of the adaptative response to NAD^+^-depletion and contributes to the establishment of resistance to NAMPT inhibition. Stable YAP downregulation strongly decreased FK866 sensitivity in TNBC and pancreatic cancer cells, in line with our recent findings in mesenchymal stem-like prostate cancer cells^17^. The resistant phenotype was not reproduced using a second short-harpin targeting YAP and transient YAP silencing resulted in only a partial reduction in FK866 sensitivity, suggesting that sustained YAP downregulation may allow the establishment of additional cellular adaptations required for resistance. Nonetheless, re-expression of constitutively active YAP5SA partially restored FK866 sensitivity in MDA shYAP cells, supporting a contribution of YAP activity to the resistant phenotype while also indicating that additional mechanisms are involved.

To explore the cellular adaptations associated with sustained YAP downregulation, we compared the proteomic profiles of MDA shYAP and RES cells, the latter of which had independently acquired resistance through prolonged FK866 exposure^12^. Interestingly, the two resistant models presented common molecular features, with a downregulation of cell-cycle, mitotic and RNA-processing programs, and an enrichment of mitochondrial-related processes. In fact, modulation of these processes has been independently associated with the cellular responses to NAD+ depletion. NAMPT inhibition has been reported to induce G2/M accumulation in different cancer models^31,32^, while YAP has been implicated in spindle-checkpoint activation through regulation of BubR1, with YAP downregulation impairing checkpoint activation following microtubule perturbation^33^. Indeed, MDA shYAP cells showed an attenuated G2/M accumulation following nocodazole treatment compared with MDA shSCR cells. The deregulation of protein translation upon FK866 exposure has also been previously described by our group, with the activation of an AMPK/EIF2A-dependent translational arrest protecting cells from the energetic stress induced by NAMPT inhibition^29^. These findings are consistent with the broader modulation of RNA-processing programs observed in both resistant models.

In parallel, mitochondrial rewiring has been previously associated with acquired resistance to FK866, with RES cells presenting increased mitochondrial mass and respiratory capacity and increased utilization of mitochondrial substrates^12^. Similarly, MDA shYAP cells showed increased mitochondrial respiration, spare respiratory capacity and mitochondrial mass. In fact, YAP/TAZ have also been associated with the regulation of mitochondrial metabolism, with their depletion promoting increased mitochondrial respiration in NF2 mutant tumors ^34^.

The concomitant modulation of cell-cycle progression, RNA/protein biosynthesis and mitochondrial function may therefore represent different components of the cellular adaptation to NAD⁺ depletion. Importantly, the NAD⁺ content and the NAD⁺/NADH ratio differ between cytoplasmic and mitochondrial compartments, meaning that these pools may also respond differently to NAD⁺ depletion^35^. It is therefore possible that the increased mitochondrial mass and function observed in MDA shYAP and RES cells contribute to the preservation of cellular NAD(H) during FK866 exposure, while decreased proliferation and protein synthesis may reduce the energetic and biosynthetic requirements of resistant cells.

Of note, the enzyme nicotinamide N-methyltransferase (NNMT) was not among the proteins quantified in our proteomics datasets, precluding the direct assessment of the YAP-NNMT axis, identified by our group as a mechanism of innate resistance to FK866 in prostate cancer ^17^.

In conclusion, our data identify YAP as a modulator of the cellular response to NAMPT inhibition in TNBC. Sustained YAP downregulation is associated with the establishment of a resistant state characterized by mitochondrial rewiring and modulation of cell-cycle and biosynthetic programs, resembling adaptations independently acquired during prolonged FK866 exposure. These findings provide a link between YAP signaling and the metabolic plasticity that allows cancer cells to tolerate NAD⁺ depletion.

## ETHICS STATEMENT

No human studies, animal studies and potentially identifiable human images or data are presented in this manuscript.

## AKNOWLEDGEMNETS

We thank Romina Belli and Daniele Peroni from DiCIBIO’s Proteomics and MS core facility at the University of Trento for performing the mass spectrometry analyses. The European Regional Development Fund (ERDF) 2014-2020 POR P.A. Trento is also acknowledged for supporting the Proteomics and MS core facility at the University of Trento.

## AUTHORS CONTRIBUTIONS

Conceptualization: AC, AP; Data curation: AC, AP Funding acquisition: AC, AP; Resources: AP; Supervision: AC, AP; Validation: AB, EF, EC, IF, VA, AC; Visualization: AC, AP; Roles/Writing - original draft: AC; Writing - review & editing: All authors.

## FUNDING

This project has received funding from the European Union’s Horizon 2020 research and innovation program, INTEGRATA, under grant agreement No 813284 and by the initiative “Dipartimenti di Eccellenza 2023-2027 (Legge 232/2016)” funded by the MUR to CIBIO Department to AP. A.C. has received funding from the European Union’s Horizon 2020 research and innovation program, under the grant agreement no. 101279563 – eN-TIME-HORIZON-MSCA-2025-PF. VA is supported by MFAG-2021 #26004 from the Associazione Italiana di Ricerca sul Cancro (AIRC).

---

**Supplementary Figure 1.**
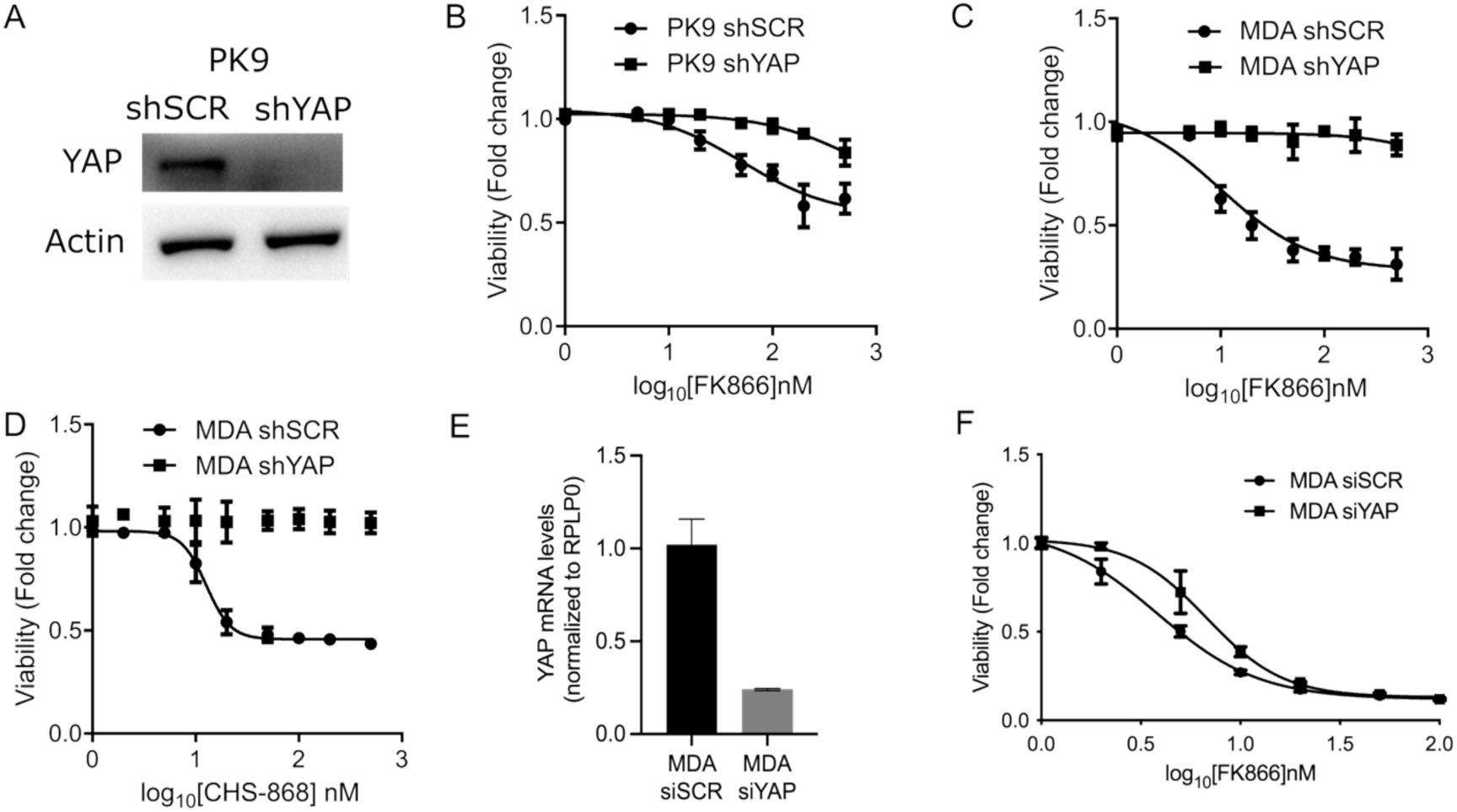
YAP downregulation decreases sensitivity to NAMPT inhibition. (A) Western blot confirming YAP downregulation in PK9 shYAP cells compared with PK9 shSCR controls. Actin was used as loading control. (B) Viability of PK9 shSCR and PK9 shYAP cells following 48 h exposure to increasing concentrations of FK866. Viability of MDA shSCR and MDA shYAP cells following 48 h exposure to increasing concentrations of FK866 (C) or CHS-828 (D), assessed by the sulforhodamine B (SRB) assay. (E) YAP expression following transient YAP silencing in MDA-MB-231 cells. (F) Viability of MDA-MB-231 cells transiently silenced for YAP following exposure to increasing concentrations of FK866. Data are presented as mean ± SEM from three independent biological replicates.

## Notes

### Competing Interest Statement

The authors have declared no competing interest.

